# The chloroplast 2-cysteine peroxiredoxin functions as thioredoxin oxidase in redox regulation of chloroplast metabolism

**DOI:** 10.1101/317107

**Authors:** Mohamad-Javad Vaseghi, Kamel Chibani, Wilena Telman, Michael Liebthal, Melanie Gerken, Sara Mareike Müller, Karl-Josef Dietz

## Abstract

Thiol-dependent redox regulation controls central processes in plant cells including photosynthesis. Thioredoxins reductively activate e.g. Calvin-Benson cycle enzymes. However the mechanism of oxidative inactivation is unknown despite its importance for efficient regulation. Here, the abundant 2-cysteine peroxiredoxin (2-CysPrx), but not its site-directed variants, mediates rapid inactivation of reductively activated fructose-1,6-bisphosphatase and NADPH-dependent malate dehydrogenase (MDH) in the presence of the proper thioredoxins. Deactivation of phosphoribulokinase and MDH was compromised in *2cysprxAB* mutants plants upon light/dark transition compared to wildtype. The decisive role of 2cysprxAB in regulating photosynthesis was evident from reoxidation kinetics of ferredoxin upon darkening of intact leaves since its half time decreased 3.5-times in *2cysprxAB*. The disadvantage of inefficient deactivation turned into an advantage in fluctuating light. The results show that the 2-CysPrx serves as electron sink in the thiol network important to oxidize reductively activated proteins and represents the missing link in the reversal of thioredoxin-dependent regulation.

## Introduction

Redox homeostasis is a fundamental property of life and intimately linked to the redox state of glutathione and protein thiols. Deviations from normal thiol redox environment with glutathione redox potentials of about -315 mM elicit alterations in gene expression, posttranscriptional processes, metabolism and also compensatory mechanisms for readjustment of the cellular redox norm (*Foyer et al., 2009*). All cells maintain a thiol redox regulatory network consisting of redox input elements, redox transmitters, redox targets, redox sensors and reactive oxygen species as final electron acceptors (*Dietz, 2008*). The chloroplast thiol network is presumably the best studied case of thiol redox regulation since thiol redox regulation of Calvin-Benson cycle (CBC) enzymes by a thiol factor was discovered as early as in 1971 (*Buchanan et al., 1971*). Later on the decisive factor was proven to be thioredoxin (Trx) (*Holmgren et al., 1977*). Genome annotations and biochemical studies elucidated the complex composition of the plastid thiol network of plastids and of plants in general (*Meyer et al., 2009*; *Buchanan, 2016*).

The plastids of *Arabidopsis thaliana* contain a set of 10 canonical Trxs (Trx-f1, -f2, -m1, -m2, -m3, -m4, -x, -y1, -y2, -z) and additional Trx-like proteins, e.g. the chloroplast drought-induced stress protein of 32 kDa (CDSP32)(*Broin and Rey, 2003*), four ACHT proteins (*Dangoor et al., 2009*), the Lilium proteins and Trx-like proteins(*Chibani et al., 2009*; *Meyer et al., 2009*). The canonical Trxs are reduced by ferredoxin (Fd)-dependent thioredoxin reductase (FTR) and themselves reduce oxidized target proteins. The FTR-pathway reduces the Trx-isoforms with distinct efficiency as recently shown by *Yoshida and Hisabori (2017)*. Well characterized Trx-targets are the CBC enzymes fructose-1,6-bisphosphatase (FBPase), NADPH-dependent glyceraldehyde-3-phosphate dehydrogenase, seduheptulose-1,7-bisphosphatase, ribulo-5-phosphate kinase (PRK) and ribulose-1,5-bisphosphate carboxylase oxygenase activase (RubisCO activase)(*Michelet et al., 2013*). The chloroplast FBPase is reduced by Trx-f with high preference (*Collin et al., 2003*). Another reductively activated target is the NADPH-dependent malate dehydrogenase (MDH) which plays a role in the export of excess reducing equivalents in photosynthesizing chloroplasts. MDH is activated if the stromal reduction potential increases (*Scheibe and Beck, 1979*) under conditions of limited availability of electron acceptors, e.g. in high light, low temperature or low CO_2_ (*Hebbelmann et al., 2012*). Activation is mediated by m-type Trxs (*Collin et al., 2003*).

Hundreds of Trx-targets and polypeptides undergoing thiol modifications have been identified in proteome studies (*Montrichard et al., 2009*). The various redox proteomics approaches employed affinity chromatography, differential gel separation and isotope coded-affinity or fluorescence-based labeling (*Mock and Dietz, 2016*). Trapping chromatography using Trx variants with mutated resolving cysteines allowed for efficient identification of Trx-targets (*Motohashi et al., 2009*). The target proteins are essentially associated with all important metabolic activities and molecular processes such as transcription, translation, turnover, defense against reactive oxygen species and also signaling pathways in the chloroplast (*Buchanan, 2016*). The enzymes are often activated upon reduction, but redox regulation of e.g. signaling components and certain enzymes involves oxidation as part of the response, e.g. in transcriptional regulation (*Dietz, 2014*; *Giesguth et al., 2015*; *Gütle et al., 2017*). The significance of controlled oxidation is most apparent if considering the metabolic transition from light-driven photosynthesis to darkness or from high to low photosynthetic active radiation. Enzymes of the CBC must be switched off upon darkening or adjusted to the new activity level in decreased light in order to prevent depletion of metabolites and de-energization of the cell (*Gütle et al., 2017*). In fact upon tenfold lowering the irradiance from e.g. 250 to 25 μmol quanta˙m^-2^ s^-1^, the CO_2_ assimilation transiently drops to CO_2_ release prior to adjustment to the new lower level. The NADPH/NADP^+^-ratio falls from 1.1 to 0.1 prior to readjustment of the previous ratio of about 1 in the lower light. Since also the ATP/ADP-ratio drops within 30 s, and thus the assimilatory power, *Prinsley et al.(1986)* concluded, that the deactivation of the enzymes occurs with slight delay, but then enables recovery of appropriate metabolite pools to proceed with carbon assimilation in the new light condition.

Reversible and rapid redox regulation requires efficient mechanisms not only for reduction but also for oxidation of target proteins. The reductive pathway via Trxs is well established, thus the open question concerns the mechanism of oxidation. The likely candidate is hydrogen peroxide as most stable and thus diffusible reactive oxygen species (ROS). The light reactions generate superoxide in the Mehler reaction, and possibly at low rates also in the reaction catalyzed by the plastid terminal oxidase (*Dietz et al., 2016*). Two molecules of superoxide are dismutated to one molecule of H_2_O_2_ and one molecule of O_2_. In particular photosynthesis-derived ROS were often considered as unavoidable side products but are now accepted drivers in the redox regulatory network and thus in regulation and signaling (*Driever et al., 2011*). ROS likely are involved in operating the regulatory thiol switches (*Groitl and Jakob, 2014*). Direct non-specific reaction of H_2_O_2_ with target proteins would lack specificity. It is also questionable that thiols of target proteins including Trxs as transmitters can compete as electron donors to H_2_O_2_ with those proteins that evolved for that purpose, the thiol peroxidases of the chloroplast (*Dietz, 2016*; *Flohé, 2016*). O_2_ appears even less suitable than H_2_O_2_ as thiol oxidant due to its relative high stability.

Thiol peroxidases are evolutionary ancient proteins that efficiently react with peroxides. The catalytic cysteinyl residue is embedded in a specific molecular environment that lowers its pK_a_ value. The high affinity with K_M_-values in the low micromolar range compensates for low turnover numbers. The Arabidopsis chloroplasts contains four peroxiredoxins (two 2-cysteine peroxiredoxins [2-CysPrxA, 2-CysPrx B], peroxiredoxin Q [PrxQ], peroxiredoxin IIE [PrxIIE]) and two glutathione peroxidase-like proteins which also function as Trx-dependent peroxidases (*Horling et al., 2003*; *Navrot et al., 2006*; *Dietz, 2016*). The 2-CysPrxA/B represent the most abundant chloroplast peroxiredoxin with about 100 μM concentration (*Peltier et al., 2006*). Antisense plants lacking the 2-CysPrx reveal disturbed photosynthesis and altered redox homeostasis (*Baier and Dietz, 1999*; *Baier et al., 2000*). Analysis of *A. thaliana* lines with T-DNA insertions established that the NADP-dependent thioredoxin reductase C (NTRC) is the predominant and efficient electron donor to 2cysprxAB (*Pulido et al., 2010*) and that the 2cysprxAB participate in detoxification of H_2_O_2_ generated in the Mehler reaction (*Awad et al., 2015*). But it is also established that 2-CysPrx accepts electrons with lower efficiency from various Trx and Trx-like proteins (*Collin et al., 2003*). Recently it was shown, that the severe growth phenotype of *ntrc*-plants can be recovered by crossing them with the *2cysprxAB* plants (*Pérez-Ruiz et al., 2017*). In the *ntrc* plants, the FBPase was insufficiently reductively activated in the light. In gel redox analysis revealed that Trx-f remained partially oxidized even in the light and thus probably was unable to activate FBPase. This effect was mostly reverted in the *ntrc/2cysprxAB*-triple mutants. The authors proposed that the lack of NTRC and the concomitant oxidation of 2cysprxAB (*Pulido et al., 2010*; *Pérez-Ruiz et al., 2017*) oxidize Trx-f. This assumption is tentatively in line with the low electron donation capacity of Trx-f to 2-CysPrx reported by *Collin et al.(2003)*.

This study aimed to dissect the hypothesis proposed in 2008 by Dietz, namely that the 2-CysPrx functions as the missing link in the thiol-disulfide redox regulatory network, as Trx-oxidase in redox regulation of chloroplast metabolism and other regulatory processes. *In vitro* experiments were designed to study the inactivation of FBPase and MDH by oxidized 2-CysPrx. The inactivation depended on the presence of Trxs with preference for specific Trxs. The hypothesis was further scrutinized *in vivo* by comparing wildtype (WT) and *2cysprxAB* for Kautzki effect, ferredoxin reoxidation kinetics, inactivation of malate dehydrogenase and ribulo-5-phosphate kinase upon darkening. The data strongly support the Trx-oxidase hypothesis and show a pre-activation of metabolism in darkness and a delayed inactivation during light-to-dark transfer. Development of a kinetic model allowed for simulating the *in vitro* observations. Finally, it will be shown that the growth inhibition phenotype of *2cysprxAB* relative to WT-plants is reversed in short fluctuating light pulses supporting the hypothesis that 2-CysPrx is needed for efficient inactivation in fluctuating light. While WT fails to use the light energy during the short light pulses, the *2cysprxAB* mutant exploits this energy given its inefficient deactivation of the redox regulated enzymes. Thus 2-CysPrx is a principle component in rapid plant light acclimation.

## Results

*A. thaliana* lacking 2-CysPrx develop with delay and show defects in photosynthesis (*Baier and Dietz, 1999*). *Figure 1* depicts chlorophyll a-fluorescence transients of WT seedlings and seedlings lacking cyclophilin 20-3, an interactor of 2-CysPrx, or 2cysprxAB when grown on solidified Murashige Skoog plus/minus sucrose. The actinic light was switched on after 15” and both wildtype and *cyp20-3* seedlings displayed the Kautsky peak of chlorophyll-a fluorescence emission, followed by a slow decline to the new steady state. In a contrasting manner, *2cysprxAB* seedlings showed a small fluorescence increase in the absence of sucrose, which was missing in the presence of sucrose. The Kautsky effect reflects the reduction of the electron transport chain in the first phase of illumination followed by reoxidation in the course of activation of NADPH- and ATP-consuming metabolic pathways, in particular the CBC and the malate valve. The lack of the fluorescence peak in *2cysprxAB* was tentatively interpreted as indication that the energy sinks and energy dissipation mechanisms such as non-photochemical or photochemical quenching activities remained active in darkness and that this pre-activation is linked to properties of 2-CysPrx.

**Figure 1.**
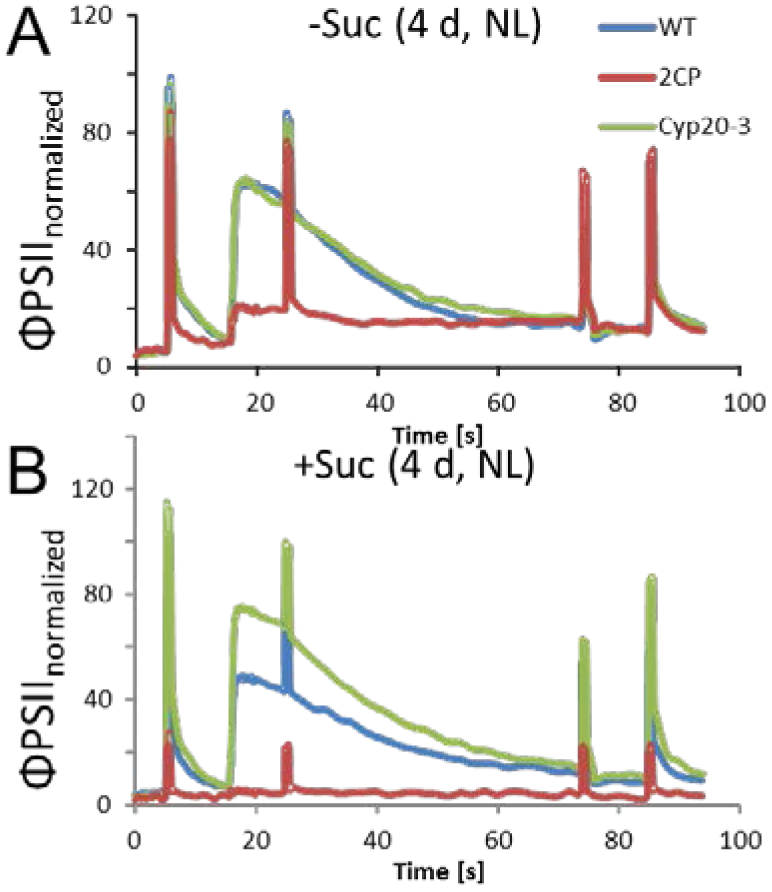
Chlorophyll-a fluorescence kinetics of 4d old wildtype, *2cysprxAB* and *cyclophilin 20-3* mutants. Seeds were placed on phytogel-solidified half strength Murashige-Skoog medium with or without 0.5% sucrose, stratified for 2 d and then grown in a growth chamber with 8h light, 16h dark at 80μmol quanta˙m^-2.^s^-1^ and 23°C. Chlorophyll a-fluorescence transients of the seedlings on the plates were imaged with the FluoroCam with the following settings: 30 min dark acclimation, measuring light on after 5s, actinic light with 65μmol quanta˙m^-2.^s^-1^ after 16.5s. Actinic light was switched off after 76.5s and the recording terminated at 94.5s. Saturating light pulses of 900 μmol quanta˙m^-2.^s^-1^ were applied after 5, 25, 75 and 85s.

Chloroplast fructose-1,6-bisphosphatase (FBPase) and NADPH-dependent malate dehydrogenase (MDH) were selected as two well-known examples for light-activated enzymes in order to address the hypothesis that 2-CysPrx affects the activation state of redox proteins. Stroma protein was obtained from isolated intact chloroplasts, and chloroplast Trxs and 2-CysPrx were generated as recombinant protein. The stroma extract was first preincubated in 1 mM dithiothreitol (DTT) for reductive activation of FBPase. The *in vitro* assay after 1:1 dilution (residual DTT concentration of 500 μM) was initiated by addition of the substrate FBP (*Figure 2*).

**Figure 2.**
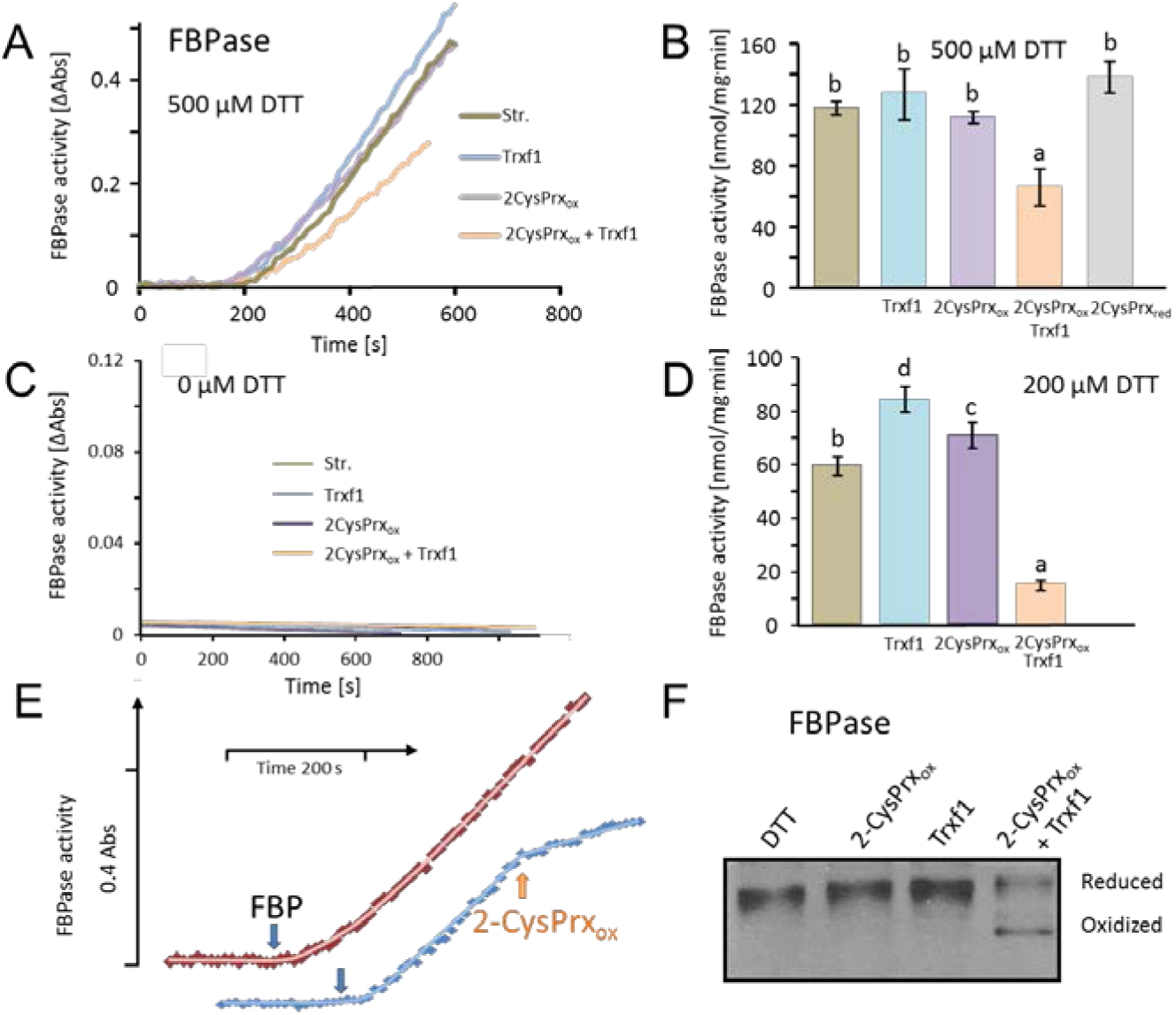
Inhibition of reductively activated FBPase by oxidized 2-CysPrxA. (A) Isolated stroma was treated with 1 mM dithiothreitol (DTT) plus/minus Trx for pre-activation and then added to the FBPase activity test with a final DTT concentration of 500 μM. Test compounds as indicated were added at t=0 s. Concentrations were: 5 μM Trx-f1, 5 μM oxidized 2-CysPrx. The enzyme assay was started by addition of 600 μM FBP at t=180 s. (B) Average activity values were calculated from the maximum slope obtained for the different treatments from independent experiments. Data are means±SD of n=3. (C) Control measurement without DTT. (D) Activity in the presence of final DTT concentration of 200 μM (pre-activation with 400 μM DTT). (E) The speed of inactivation was documented by addition of 5 μM 2-CysPrxAox in the presence of Trx f1 at the time point as indicated after establishing the linear rate of FBP hydrolysis. The FBPase activity was rapidly inhibited after addition of 2-CysPrxAox. (E) Redox state of FBPase in the assay shown in *Figure 1A* after labeling the reduced thiols with mPEG_24_. Migration of the reduced 2-CysPrx was slowed down and was abundant in the samples treated with DTT, DTT plus 2-CysPrxox or DTT plus Trx-f1 (upper band), but was strongly decreased after addition of the combination of 2-CysPrxox and Trx-f1. Same results were seen in several experiments.

The stromal FBPase efficiently converted FBP to Fru-6-P as seen from the time-dependent increase in absorption. Addition of Trx-f1 slightly increased the activity of FBPase indicating that activation had not been fully achieved during the preincubation, so that Trx-f1 in the presence of the residual 500 μM DTT was able to further reduce and activate the stromal FBPase. The activity of FBPase was unchanged upon addition of oxidized 2-CysPrx in the absence of Trx. However if the complete test was reconstituted with reduced stroma, Trx-f1 and oxidized 2-CysPrx, FBPase was significantly inhibited. Apparently, Trx-f1 mediated the oxidative inactivation of FBPase by transferring electrons from reduced FBPase to 2-CysPrx_ox_. As an additional control reduced 2-CysPrx was added to reduced stroma and established the same activity as with Trx-f1 alone (*Figure 2A, B*). Supplementary Figure 1

Activity was not seen in the absence of DTT (*Figure 2C*). Overall activation was lower if the pre-activation was performed with 400 μM instead of 1 mM DTT (*Figure 2D*). Under these conditions, addition of reduced Trx-f1 further activated the FBPase. Addition of 2-CysPrx_ox_ to reduced stroma again had no effect on FBPase activity, but the combination of reduced stroma, Trx-f1 and 2-CysPrx_ox_ led to 76% inhibition. The inhibition was rapidly achieved as revealed when 2-CysPrx_ox_ was added to the ongoing enzyme reaction (*Figure 2E*). In parallel, the reduced form of FBPase which runs as upper band in *Figure 2F* due to incorporation of 2 molecules of mPEG_24_ per molecule FBPase only disappeared in the fully reconstituted assay with reduced stroma, Trx-f1 and 2-CysPrx_ox_.

The efficiency of inactivation depended on Trx-f1 amounts (*Figure 3A*). Thus inhibition was low in the presence of 0.625 μM Trx-f1, while a strong decrease was detected in 5 μM Trx-f1,revealing saturation with increasing Trx-f1 concentration. Comparison of inactivation in the presence of Trx-f1, -m1, -m4, -x and CDSP32 revealed exclusive preference for Trx-f1 (*Supplementary Figure 1*).

**Figure 3.**
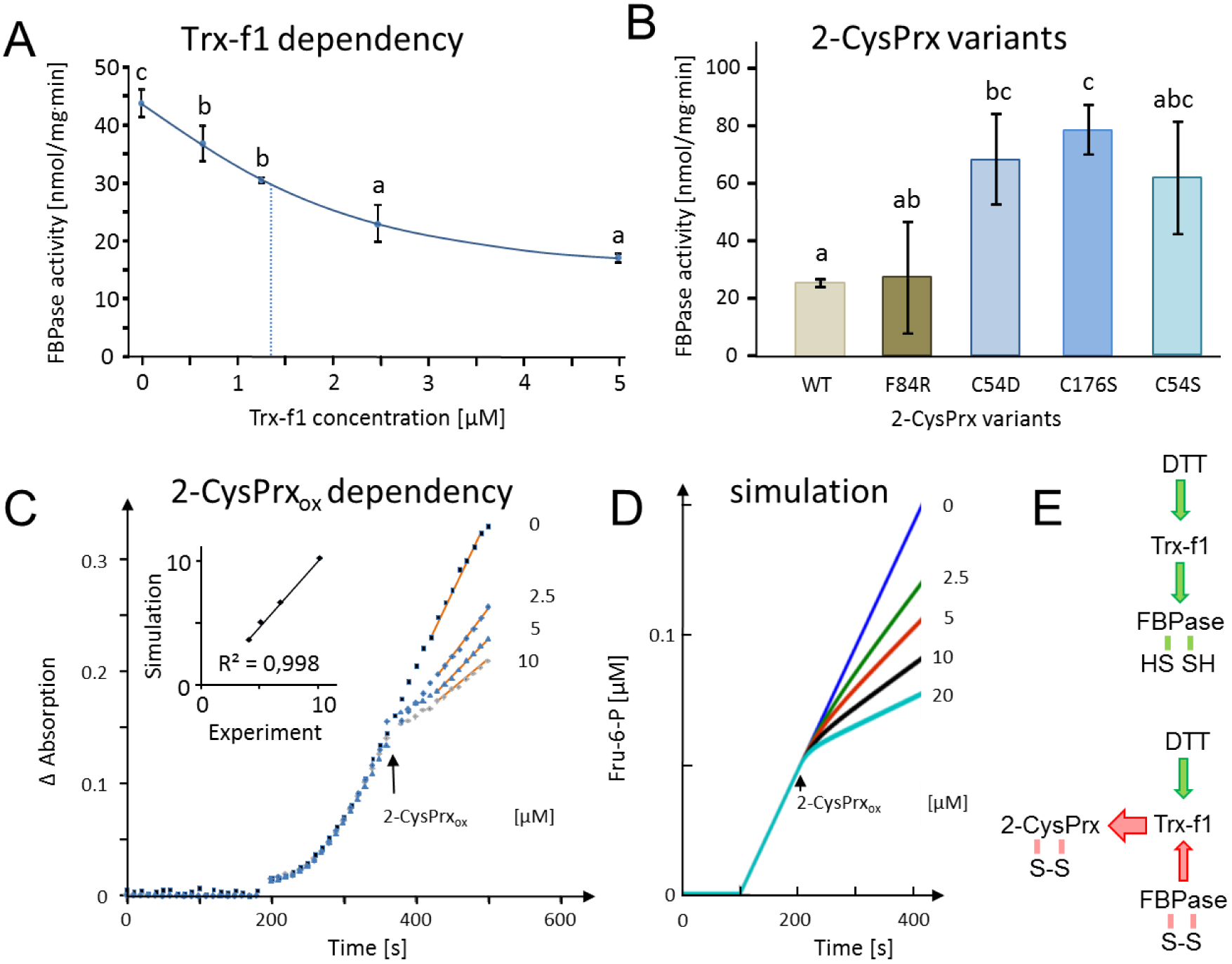
Dependency of FBPase inactivation on Trx-f1 concentration and functionality of the 2-CysPrxA, and mathematical simulation of the *in vitro* assay by kinetic modeling. (A) Dependency of FBPase activity on Trx-f1 concentration. At t=0 min, Trx-f1 was added at concentration between 0 and 5 μM. The FBPase activity test was performed as in Figure 1A. Data are means±SD of n=3. (B) Requirement of the functional thiol peroxidase for FBPase inhibition. The test contained 5 μM Trx-f1, final concentration of DTT was 500 μM and the 2-CysPrx variants were added at 5 μM concentration. The WT form of 2-CysPrxox inhibited the FBPase by 75%, as did the F84R variant which is compromised in oligomerization. The variants devoid of the peroxidatic Cys in C54S and C54D, as well as the variant lacking the resolving Cys in C176S were ineffective in inhibiting the reductively activated FBPase. (C) Effect of adding different final 2-CysPrxox concentrations to the ongoing FBPase activity test. Conditions were as described in Figure 1A. The traces are means of n=3 determinations. (D) Mathematical simulation of the concentration-dependent effect of 2-CysPrxox on FBPase activity. Addition of 0, 2.5, 5 and 10 μM, and also 20 μM 2-CysPrx_ox_ was simulated (the effect of 20 μM 2-CysPrxox could not be tested experimentally). The relative slopes after addition of 2-CysPrxox in the experimental and theoretical analyses were plotted and gave a linear dependency with a regression coefficient of 0.998 (inset in *Figure 2C*). (E) Schematics of the measured and simulated pathways before (upper) and after addition of 2-CysPrxox (lower scheme). Green arrows represent reductive activation, red arrows oxidative inactivation.

CysPrx employs two Cys-residues, the peroxidatic Cys54 and the resolving Cys176, in the detoxification reaction of peroxides such as H_2_O_2_, alkylhydroperoxides and peroxinitrite. The dependency of the Trx oxidase activity of the 2-CysPrx on the presence of the catalytical thiols was tested by using site-directed mutated variants of 2-CysPrx (*König, 2013*)(*Figure 3B*). The variants C54S, C176S and also the hyperoxidation mimicking C54D were unable to inactivate FBPase, thus both thiols are needed. In a converse manner, the variant F84R which is compromised in its decamerization ability (*König, 2013*) but still acts as thiol peroxidase efficiently inactivated the FBPase in the complete inactivation assay.

The *in vitro* FBPase activity was simulated by use of a kinetic mathematical model (*Figure 3E*; *Supplementary Tables 1-4*; *Supplementary Figure 2*) and the results compared with an experiment where oxidized 2-CysPrx was added to the enzyme assay at different concentrations (*Figure 3C, D*). The addition of 2.5, 5 or 10 μM 2-CysPrx_ox_ progressively inhibited the turnover of FBP to Fru-6-P as indicated by absorption change. The simulation (*Figure 3E*) started with fully activated FBPase in the presence of 5 μM Trx-f1. Addition of 2-CysPrx_ox_ at 2.5, 5, 10 or 20 μM led to rapid partial and concentrations-dependent inhibition of FBPase with high fit to the empirical data.

The NADPH-dependent malate dehydrogenase (MDH) is another established target of redox regulation in the chloroplast. MDH is activated by Trx-m if the reduction potential of the stroma turns highly negative e.g. in excess light. MDH activity was measured *in vitro* under similar conditions as the FBPase described above. Reductive pre-activation of MDH enabled the conversion of oxaloacetic acid to malate with concomitant oxidation of NADPH (*Figure 4A*). Addition of 2-CysPrx_ox_ had no influence on MDH activity, showing full activation by DTT. The supplementation of the assay with various Trxs during pre-incubation enabled the inactivation of MDH after addition of 2-CysPrx_ox_. Thus, Trx-m1 mediated complete MDH inhibition by 2-CysPrx_ox_. The chloroplast drought specific protein CDSP32 ranked second among the tested Trxs (*Figure 4B*). Thus the overall order of inhibition efficiency was Trx-m1>CDSP32>Trx-f1>Trx-x>Trx-m4.

**Figure 4.**
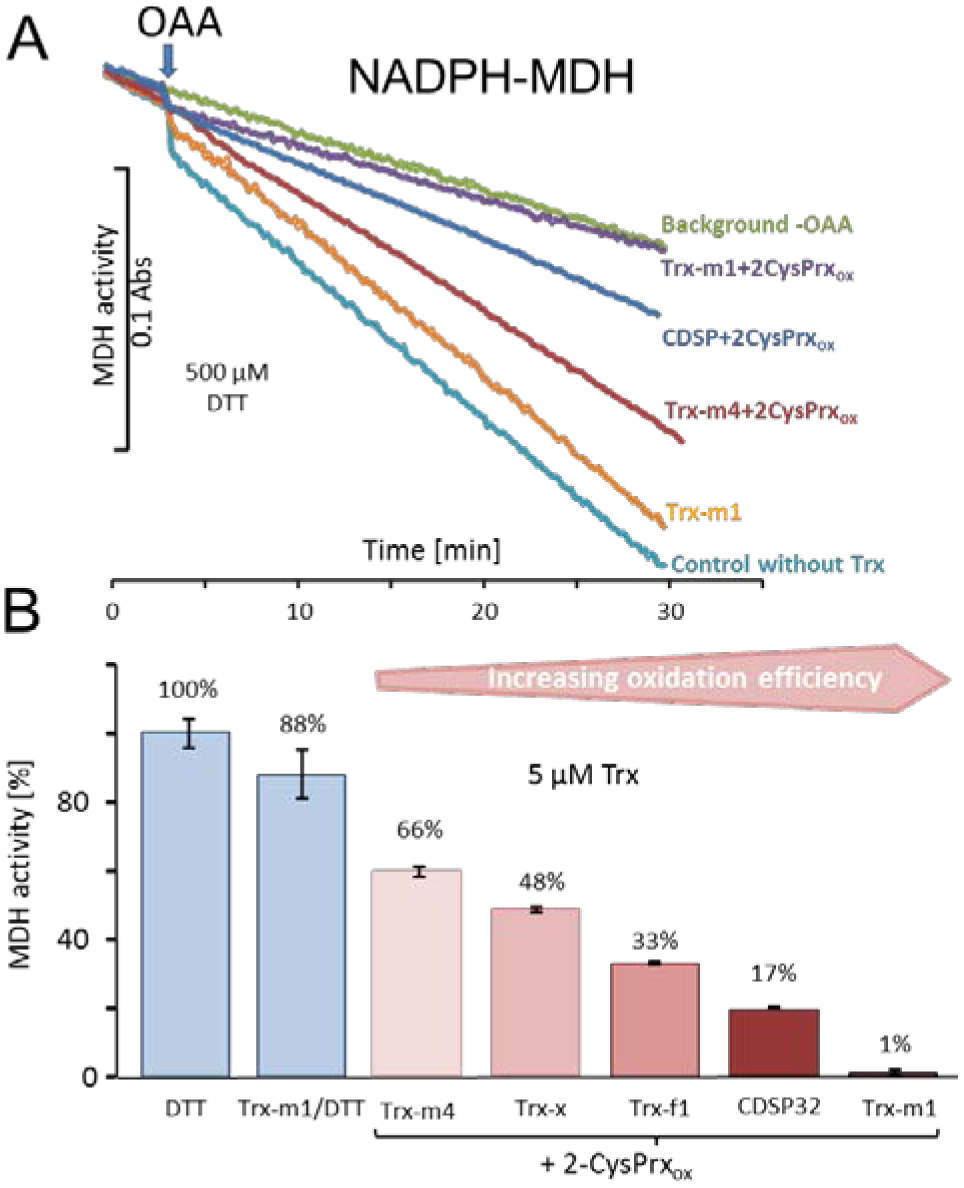
Trx-dependent inactivation of MDH by oxidized 2-CysPrx. (A) Spectrophotometric recording of MDH activity after addition of 2 mM oxaloacetic acid (OAA). The background was determined without addition of OAA. Addition of 10 μM Trx-m1 did not alter the turnover rate. Supplementation with combinations of different Trxs and 2-CysPrxox inhibited the MDH to a varying degree. (B) Average inhibition of MDH activity in experiments as shown in (A). The numbers above the bars indicate the percentage of inhibition realized by the various Trxs. Data are means±SD of n=3.

Inactivation of reductively activated proteins by oxidation is considered to be essential in photosynthesis upon lowering the photon flux density and darkening. The inhibition of CBC enzymes, ATP synthase and MDH prevents depletion of metabolites, suppresses futile cycles and long-lasting de-energization. MDH activity was measured in leaf extracts of high light-illuminated WT plants and the *2cysprxAB* insertion mutant in a time course following darkening (*Figure 5A*). MDH was rapidly inactivated in WT, while inhibition was delayed in *2cysprxAB*. The inactivation state was significantly different at 10 s after darkening of WT and *2cysprxAB*.

**Figure 5.**
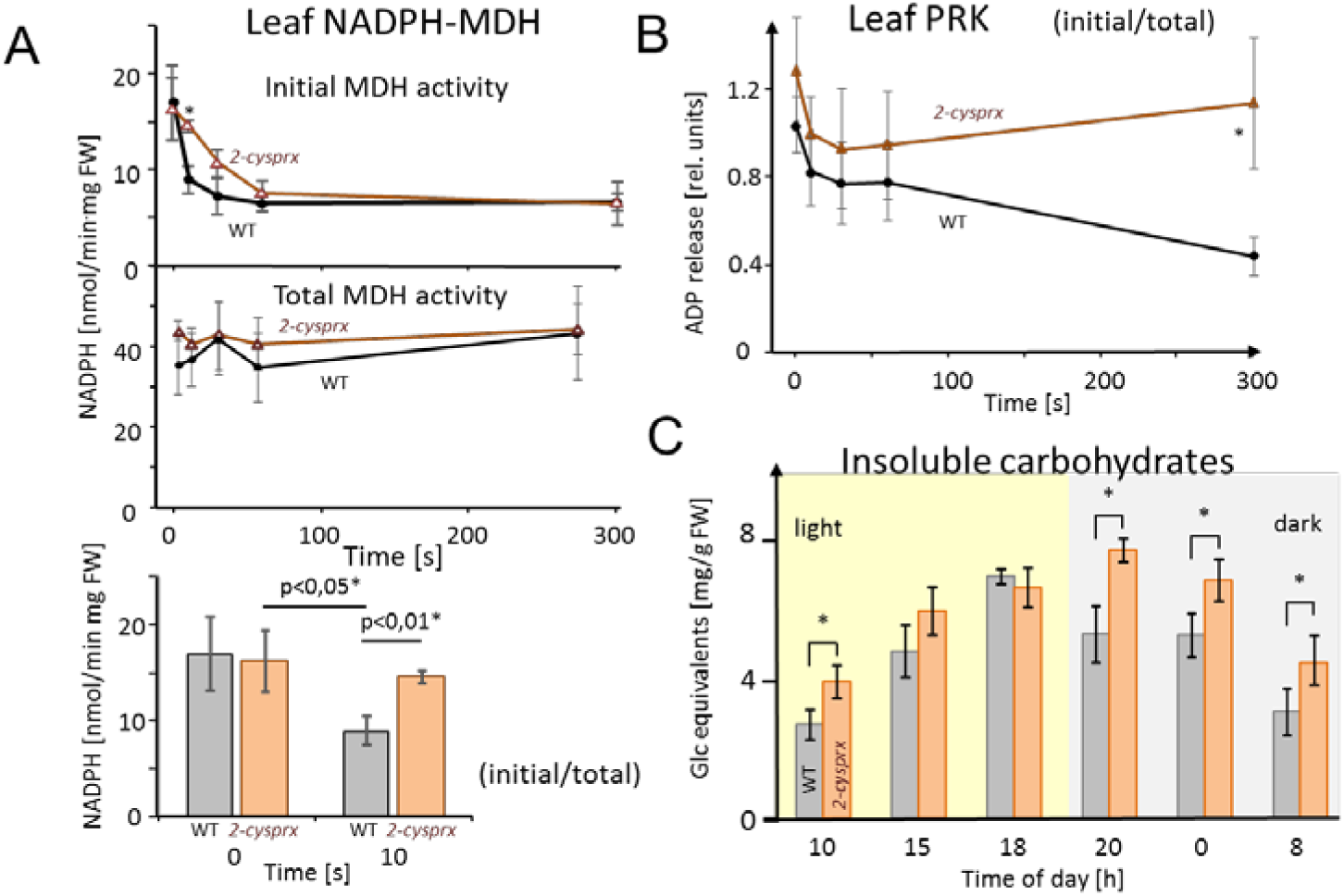
Inhibition of NADPH-MDH and phosphoribulokinase in leaves upon light-dark transitions, and non-soluble sugar contents in WT and 2cysprxAB during a 24 h day-night cycle. (A) MDH activity during a light-dark transition: WT and 2cysprxAB plants were exposed to 650 μmol quanta/m^-2^s^-1^ for 30 min and then darkened at t=0 s. Proteins were rapidly extracted prior to darkening and after different time points in the dark as indicated. Initial (upper figure) and total MDH activity (lower figure) after full activation with 500 μM DTT were determined. Inactivation of MDH was slightly delayed in 2cysprxAB than in WT. This is shown for the time point 10 s. Data are means±SD from n=3 determination. (B) Phosphoribulokinase (PRK) activity during a light-dark transition plotted as ratio of initial to total activity. WT and 2cysprxAB plants were illuminated with 650 μmol quanta/m^-2^s^-1^ for 30 min and darkened as above. Initial and total PRK activities were determined and are plotted as ratio. Initial inactivation was insignificantly delayed in 2cysprxAB, but while PRX in WT continued to be inactivated until 5 min, the PRK activity essentially remained unaltered and became significantly different at t=300 s. (C) Changes in insoluble carbohydrates during a 24 h day-night cycle. Leaves of WT and 2cysprxAB plants were harvested at 10 o’clock (1 h after start of light phase), 15, 18, 20 (1 h after end of light phase), 0 and 8 (1 h before end of dark phase). Insoluble acid hydrolysable carbohydrates were quantified in the washed sediment of leaf homogenates using the Anthron reagent following 1 h boiling in sulfuric acid. Data are means±SD of n=3.

Ribulose-5-phosphate kinase (PRK) was chosen as another reductively activated CBC enzyme which catalyzes the committed step of ribulose-1,5-bisphosphate generation for RuBP carboxylation/oxygenation. The time course of PRK activity was followed during a light-dark transition and revealed deactivation in WT with progression of darkness while the activity remained unchanged in *2cysprxAB* (*Figure 5B*).

To address the more global effect of lacking 2-CysPrx, insoluble acid hydrolysable carbohydrates were measured during a day time course (*Figure 5C*). The amounts of insoluble carbohydrates increased 2.6-fold during the light phase and declined during the night in WT. Carbohydrate amount was 40% higher in *2cysprxAB* at the end of the dark phase and reached a similar amount at the end of the light phase as in WT. Mobilization in the dark phase started with a delay.

Ferredoxin is the central electron distributor transferring electrons from the photosynthetic electron transport chain (PET) to consuming metabolic reactions such as Fd-dependent NADPH reductase, Fd-nitrite reductase, Fd-sulfite reductase, Fd-Trx-reductase and others. The re-oxidation rate of Fd was determined in leaves from WT (*Figure 6A*) and *2cysprxAB* (*Figure 6B*) using the near infrared kinetic LED spectrometer (NIR KLAS 100). Dark adapted leaves were illuminated with a short 1.5 s light pulse of 162 μmol quanta m^-2^s^-1^ and then darkened again. During this short light period the Fd pool was photoreduced. The reoxidation of the Fd pool was followed as difference between the NIR absorption of the leaf at 785 and 840 nm (*Klughammer and Schreiber, 2016*). In WT plants, reoxidation occurred with a half-life time of 416±81 ms, while Fd reoxidized with a half-life time t_50_ of 120±35 ms in 2cysprxAB (n=5, m±SD) resulting in a very stable increase in oxidation rate by a factor of 3.5 if 2-CysPrx is missing. This decrease in t_50_ indicates that the oxidative inactivation of electron-consuming reactions was effective in WT but markedly delayed in plants lacking 2cysprxAB.

**Figure 6.**
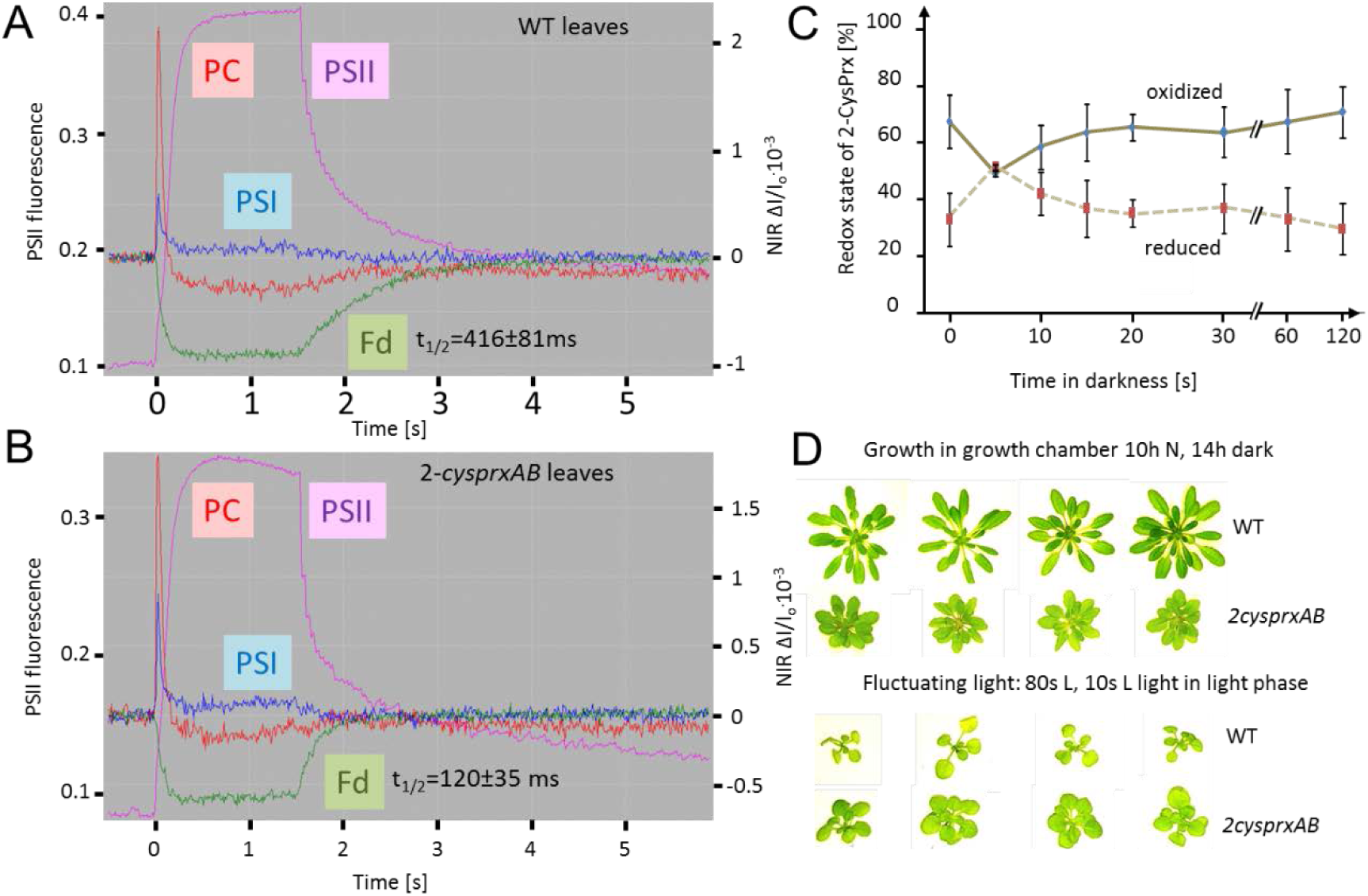
Reversal of light induced changes of photosynthetic parameters. (A) WT and (B) *2-cysprxAB* plants were acclimated to darkness. Fluorescence and NIR absorption changes were recorded with the NIR-KLAS-100. Chlorophyll-a fluorescence from photosystem II (PSII, violet trace) (left ordinate) and redox changes from photosystem I (PSI, blue), plastocyanin (PC, red) and ferredoxin (Fd, green) (right ordinate) were measured during a fast kinetics analysis consisting of 1.5 s illumination with 162 μmol quanta m^-2^ s^-1^ followed by darkening. The analysis of the fluorescence and absorption decay showed slower half-life times for WT than for *2cysprxAB* plants. The figure shows recordings typical for all analysis with n=5 measurements. (C) Redox state of the 2-CysPrx ex vivo during a light dark transition. WT plants were acclimated to 650 μmol quanta.m^-2^s^-1^ for 30 min (t=0 s) and then darkened. Samples were harvested (t=5 s-120 s) and blocked with 100 mM N-ethylmaleimide, separated by SDS-PAGE, immunodecorated with anti-2-CysPrx antibody and visualized by luminescence detection. The band intensities on shortly exposed films were quantified by ImageJ. Data are means±SD of n=4 experiments. (D) WT and *2cysprxAB* plants were grown in normal light (NL) for 42d (upper row) or for 14d in N light followed by 21d in fluctuating light with 80” L /10” H during the 10h light phase (lower row). Growth of *2cysprxAB* was inhibited in NL relative to WT, but the growth performance was reversed in fluctuating light. Four plants are shown for each growth condition and genotype.

The redox state of 2-CysPrx was assessed during the light-dark-transition. In non-reducing gel separations of N-ethylmaleimide-blocked samples, the 2-CysPrx monomer at about 22 kDa corresponds to the reduced fraction, the dimer at 44 kDa to the fully or partially oxidized form. In the light, only about 40% of total 2-CysPrx was reduced. This fraction increased upon darkening and then decreased during extended darkness (*Figure 6C*).

A considerable portion of the 2-CysPrx was oxidized both in the light and in the dark with transient variation during light transitions which might indicate the involvement in oxidation of Trx and Trx-dependent targets.

The previous data are in line with less efficient inactivation of metabolism in 2cysprxAB compared to WT. This hypothesis was further scrutinized by a growth experiment with WT and *2cysprxAB* plants assuming that the slow inactivation of metabolism in *2cysprxAB* might be advantageous under certain fluctuating light conditions with short light pulses. WT is known to outperform *2cysprxAB* (*Figure 6D*) under normal growth conditions with continuous light during the day. This advantage was challenged under fluctuating light. While WT grew better in continuous short day illumination, this changed when WT and *2cysprxAB* grew in 80 s L/10 s H. Here growth of *2cysprxAB* slightly surpassed growth of WT (*Figure 6D*).

**Figure 7.**
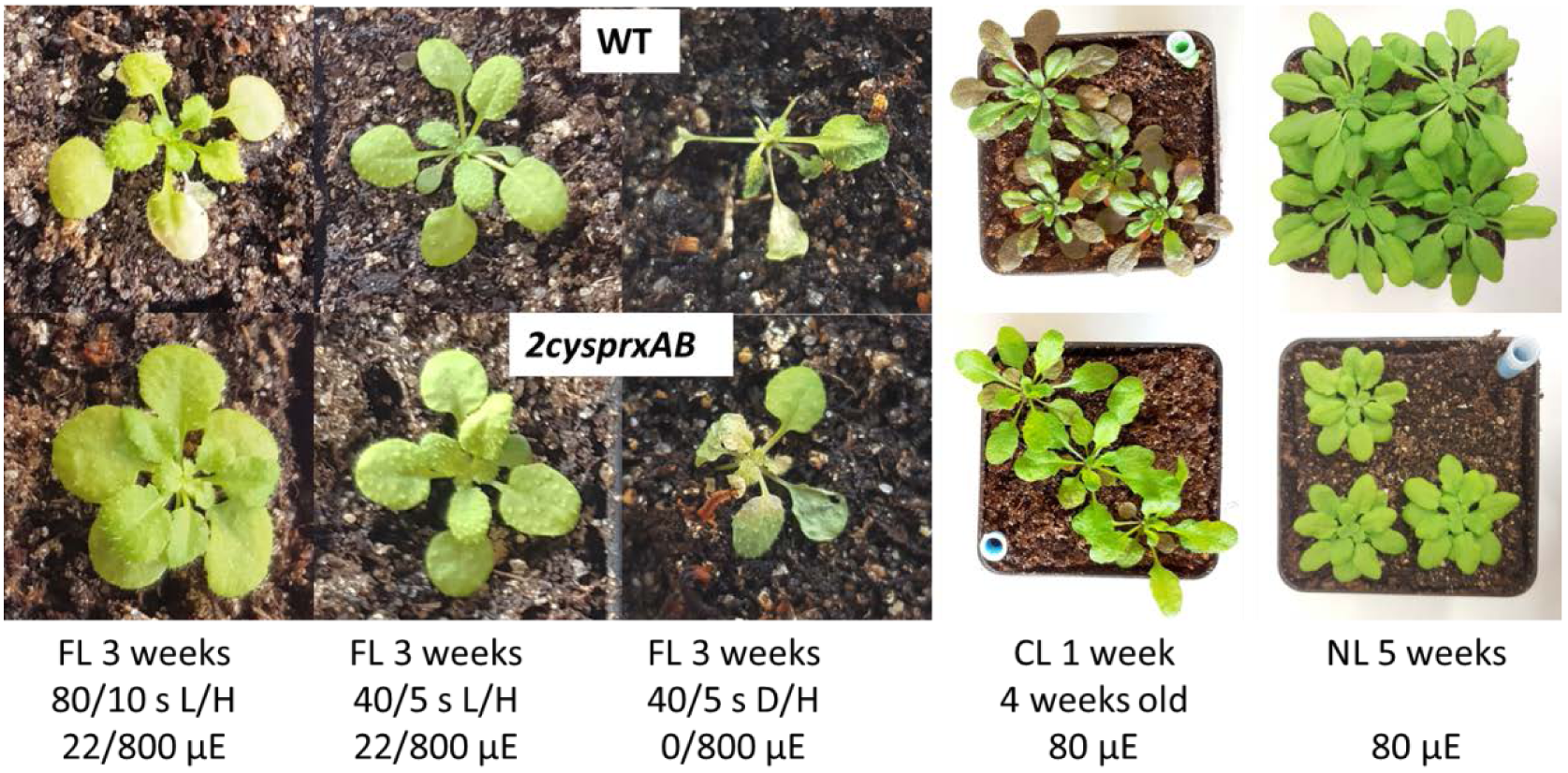
Growth phenotype of WT and *2cysprxAB* in different light regimes. WT (upper image) and 2cysprxAB (lower image) were grown in five different light regimes as indicated, namely from left to right: (i) fluctuating light (FL) for 3 weeks consisting of 80”/10” L/H with 22/800 μmol quanta.m^-2^s^-1^ (as in *Figure 5D*). The daily light cycle was maintained at 14h dark phase/10h light phase. (ii) 3 week FL with 40”/5” L/H with 22/800 μmol quanta.m^-2^s^-1^. (iii) 3 week FL with 40”/5” darkness/H with 0/800 μmol quanta.m^-2^s^-1^. (iv) Growth in continuous light without dark phase for 1 week after 4 week of growth in the growth chamber. (v) 5 week old plants grown in the growth chamber with 10h light/14h dark phase at 80 μmol quanta.m^-2^s^-1^. These growth experiments were independently performed three times with same result. Growth data are given in *Table 1*.

**Table 1.** Effect of fluctuating light on rosette fresh weight of *2cysprxAB* and WT. Plants were grown in fluctuating light for 3 weeks. The program established the following cycles: 40s L/5s H, 80s L/10s H, 120s L/15s H; H corresponds to 800 μmol quanta m^-2^ s^.1^, L to 22 μmol quanta m^-2^ s^.1^ and normal growth light (80-100 μmol quanta m^-2^ s^.1^). Plants were harvested and analyzed for biomass accumulation depending on different light/dark schemes.

| light cycle | WT [mg] | *2cysprxAB* [mg] | Ratio<br>WT/*2cysprxAB* |
| --- | --- | --- | --- |
| constant | 372±74 | 117±20 | 3.19 |
| 40s/5s | 26±5 | 22±4 | 1.21 |
| 80s/10s | 21±6 | 24±5 | 0.87 |
| 120s/15s | 14±3 | 11±2 | 1.34 |

The difference was less pronounced in 40’ L, 5’ H-cycles and in 40 s darkness, 5 s H, where the WT showed severe damage after 3 weeks, while the damage was less in *2cysprxAB* (*Figure 7*). Another difference was seen in continuous light without dark period. Rosette growth was rather similar: However, WT plants accumulated anthocyanins in older leaves while *2cysprxAB* remained green in all leaves of the rosette, indicating additional effects of the lack of 2-CysPrx on plant development and metabolism apart from being involved in the adjustment of metabolism during light shifts and light/dark transitions.

## Discussion

The Trx-dependent reductive pathway of redox regulation participates in the control of most cellular processes (*Buchanan, 2016*). However an open question concerns the mechanism of Trx re-oxidation which is employed to inactivate once reduced target proteins. Only the controlled interplay of reduction and oxidation enables efficient and fine-tuned regulation. The central role of H_2_O_2_ in tuning development and acclimation is generally accepted, but by which mechanism comes H_2_O_2_ into play? The properties of the oxidant(s) must meet several criteria in particular concerning the specificity, the thermodynamics including midpoint redox potential and sufficient amounts of oxidant buffer. The results from this study show that the chloroplast 2-CysPrx functions as peroxide-dependent thioredoxin oxidase. In the following we discuss the three criteria and then the more general implications.

### Specificity of oxidation

The chloroplast thiol network is hierarchically structured. High priority for activation and likely also for inactivation has the γ-subunit of the F-ATP synthase (*Gütle et al., 2017*). Tight control is needed to initiate ATP synthesis as soon as the light-driven proton motive force (PMF) is generated, and also to inhibit the ATP synthase upon lowering or extinguishing the incident light in order to avoid ATP hydrolysis upon dissipation of the PMF (*Mills and Mitchell, 1982*). These authors proposed the existence of an oxidation system in the stroma that oxidizes the γ-subunit of the F-ATP synthase in the dark. Thiol-controlled activation of CBC enzymes such as FBPase, SBPase and PRK is of second priority in order to initiate carbon assimilation and high rate energy drainage. Thiol-dependent activation of NADPH-malate dehydrogenase comes third since the malate valve should only drain reductive power to the cytosol if the CBC fails to consume the available energy provided by the PET (*Backhausen et al., 1994*). It is accepted that this specificity of reductive activation is realized by the complex Trx system of the chloroplast (*Thormählen et al., 2017*).

In contrast to activation, the inactivation has scarcely been studied. Direct oxidation of the target proteins by H_2_O_2_ neither can provide specificity nor adequate prioritization. The Trxs which are instrumental in activating the targets and in addition the Trx-like proteins like ACHT1-4 provide a platform which could assist in mediating specificity and prioritization. This is seen in the results from the *in vitro* inactivation assays. Among the tested Trxs, FBPase was efficiently inactivated by Trx-f1 only. In case of MDH, Trx-m1 was most effective, but CDSP32 and Trx-f1 and Trx-m4 also inactivated MDH with lower efficiency. Apparently, the efficiency for inactivation of FBPase and MDH revealed the expected Trx specificity. Thus complete switch off of MDH was achieved by the combination of Trx-m1 and 2-CysPrx_ox_, while the FBPase was not entirely inhibited even in the presence of the preferred Trx-f1 under the chosen conditions. The simulation of the enzyme assay in a mathematical model showed that the 2-CysPrx-dependent inactivation of FBPase was in line with the reported kinetic constants and redox potentials.

Specificity in reoxidation could also arise from the five different thiol peroxidases targeted to the plastids, namely in addition to 2-CysPrx PrxIIE, PrxQ and glutathione peroxidase-like 1 and 7 (*Dietz, 2016*). The combinatorial network of about 20 plastidial Trxs and Trx-like proteins with 5 thiol peroxidases provide a framework for tuned activation and inactivation dependent on interaction ability, concentration and redox potentials.

### Thermodynamics of oxidation

The redox midpoint potential of 2-CysPrx was reported with -315 mV (König et al., 2002), while the E’_M_ of NADPH-MDH from Sorghum and of spinach FBPase were determined with -330 mV, and -290 mV for spinach PRK (*Hirasawa et al., 1999*; *Hirasawa et al., 2000*). The E’_M_ of spinach and pea Trx-f1 with -290 mV was less negative than the E’_M_ of spinach Trx-m with -300 mV (*Hirasawa et al., 1999*). Thus oxidized 2-CysPrx is thermodynamically able to withdraw electrons from MDH and FBPase through Trx. This electron drainage should be less efficient in the case of PRK because of its less negative E’_M_. This was confirmed when estimating the initial vs. total PRK activity in leaves during a light-off experiment. PRK deactivated slowly in WT, while its activity remained unchanged in *2cysprxAB*. Thus additional mechanisms must participate in the regulation of PRK.

The oxidized CP12 protein with disordered regions functions as chaperone to assemble an inactive supramolecular complex of glyceraldehyde-3-phosphate dehydrogenase and PRK (*Graciet et al., 2003*). In the regulatory scenario of the chloroplast, the photosynthetic light reaction feeds electrons via Fd into the Trx-system. The high free energy change in electron transfer from Fd to Trx and target proteins efficiently drives their reductive activation. That is why in chloroplasts Trx reduction in the light is kinetically linked to Fd and not NADPH. Upon a drop in light intensity, the photosynthetic electron transfer to Fd ceases and Trxs are oxidized by drainage of electrons to 2cysprxAB. The transfer efficiency is not uniform, but shows significant specificity (i) through the Fd-dependent Trx reductase (*Yoshida and Hisabori, 2017*), (ii) between Trxs and targets and (iii) between Trxs and 2cysprxAB (this paper) (*Collin et al., 2003*). The efficiency of oxidation could be higher if the E’_M_ of 2-CysPrx were higher (less negative). Since the oxidation reaction persists in the light and is counteracted by NADPH/NTRC (*Pérez-Ruiz et al., 2017*), reduction and oxidation of targets essentially comprises a futile cycle which should not consume too much energy. Thus the evolutionary selection of a rather negative E’_M_ for 2-CysPrx might be a characteristic which allows achieving this goal. But in addition, one should keep in mind that 2-CysPrx undergoes conformational changes and possibly other posttranslational modifications which might affect its E’_M_ which should be explored in future work.

### The oxidant buffer pool size

The two highly identical isoforms of 2-CysPrxA and B represent very abundant proteins in the Arabidopsis stroma with a total concentration of more than 100 μM (*König et al., 2002*). A major function of 2-CysPrx is seen in its thiol peroxidase activity, participating in the ascorbate-independent water-water cycle and detoxification of Mehler reaction-derived H_2_O_2_ (*Dietz et al., 2006*; *Awad et al., 2015*) and, thereby, keeping the H_2_O_2_ concentration low. However, for most efficient thiol peroxidase function, the 2-CysPrx should ideally be fully or highly reduced. Surprisingly, the 2-CysPrx is highly oxidized under most conditions with less than 50% share of 2-CysPrx in the reduced form (*Pulido et al., 2010*) (*Figure 6C*). The rate of PET-dependent release of superoxide and subsequent H_2_O_2_ is considered to be low under growth conditions and may have regulatory function even under moderate stress (*Driever and Baker, 2011*). The oxidized fraction of ≥50% of the 2-CysPrx corresponds to a resting pool of 30-50 μM disulfide available for drainage of electrons from the Trx/Trx-like proteins and target pools. The 2-CysPrx redox state in the steady state underestimates the capacity for oxidation of target protein since the thiol-disulfide redox network is a dynamic flux system where H_2_O_2_ oxidizes and NTRC and Trx reduce the 2-CysPrx. In this context it is interesting that PrxIIE, the other soluble peroxiredoxin of the stroma has a midpoint redox potential of -288 mV (*Horling et al., 2003*), and thus could participate in oxidizing target proteins with less negative thiol redox potential.

### The regulatory context of metabolism

The effect of the 2-CysPrx_ox_/Trx-system on certain CBC enzymes and the malate valve is evident from both our *in vitro* analyses and *in vivo* data including the reoxidation kinetics of Fd in leaves upon darkening. Interestingly, the regulatory impact of 2-CysPrx on the metabolic state of the chloroplast goes far beyond carbon fixation and export of excess reducing power. The diurnal carbohydrate dynamics in *2cysprxAB* was disturbed. The carbohydrate amount in *2cysprxAB* reached its maximum in the early night phase and exceeded that of WT during the night and in the early day phase. Starch synthesis and degradation are subjected to redox regulation (*Santelia et al., 2015*). Trx-f1 contributes to activation of ADPglucose pyrophosphorylase. Plants lacking Trx-f1 accumulate less starch and more soluble sugars in the light (*Thormählen et al., 2013*). The changes in diurnal carbohydrate pattern indicate that starch degradation is impaired in *2cysprxAB*. Enzymes involved in starch degradation also exhibit redox sensitive and regulatory thiols. However degradation enzymes are reported to be more active in the reduced than oxidized state (*Santelia et al., 2015*), thus the exact role of 2-CysPrx in tuning the activity of starch turnover needs future analysis.

Other metabolic changes concern the accumulation of the aromatic amino acids phenylalanine and tryptophan which decreased in *2cysprxAB*, and the inhibited ability of *2cysprxAB* to synthesize anthocyanins in high light (*Awad et al., 2015*; *Müller et al.., 2017*). Phenylalanine synthesis proceeds in the chloroplast via arogenate (*Jung et al., 1986*) and phenylalanine availability determines anthocyanin synthesis (*Chen et al., 2016*). Redox-dependent regulation of the involved metabolic pathways still needs to be explored. However the *2cysprxAB* phenotype of compromised anthocyanin accumulation resembles the phenotype observed in ascorbate deficient mutants which was linked to regulation of genes involved in anthocyanin synthesis (*Page et al., 2012*).

The contribution of 2-CysPrx to redox regulation appears particularly important in low light, in darkness and in fluctuating light. Thus the phenotype of inhibited rosette growth with more round shaped leaves and short petioles (*Pulido et al., 2010*; *Awad et al., 2015*) was mostly reversed to WT phenotype in continuous light and high light. Fine-tuning of metabolic activities by 2-CysPrx-dependent oxidation counteracts reductive activation and only the proper balance between reduction and oxidation allows for optimized acclimation to the prevailing environmental conditions. High rate photosynthesis in high light requires little oxidative drainage of regulatory electrons. At lower light intensities the ratio of oxidation to reduction should increase for down-regulation of enzyme activities. If this oxidative counterbalance by 2-CysPrx is missing, enzymes maintain higher activity than needed, and this causes metabolic imbalances, sustains futile cycles and compromises growth performance of *2cysprxAB*.

The disadvantage of lacking 2-CysPrx converts to an advantage if the fluctuating light regime adopts a particular cycle frequency where deactivation by 2-CysPrx_ox_ in WT proceeds too fast to exploit the short subsequent light phase. In a converse manner, the delayed regulation in *2cysprxAB* allows for exploiting the short high light phase for carbon assimilation. This scenario explains, why *2cysprxAB* plants outperform WT plants in the 80 s L and 10 s H-light cycle.

The kinetic model of the *in vitro* enzyme test simulated the effect of oxidized 2-CysPrx on FBPase activity extremely well. Thus the correlation between the experimental data and the simulated data after supplementation with 0, 2.5, 5 and 10 μM 2-CysPrx_ox_ gave a linear dependency with a regression coefficient of 0.998. In a converse manner, the simulation of the light-dark transition did not realize full inhibition of FBPase in the simulated night. In this context it is important that other biochemical parameters such as FBP- and Fru-6-P-levels, Mg^2+^, Ca^2+^ and pH affect the redox sensitivity of FBPase (*Chardot and Meunier, 1991*). FBP is present at very low amounts in chloroplasts of darkened leaves (*Dietz and Heber, 1984*). The drop in FBP concentration is suggested to ease FBPase deactivation. Ca^2+^ fluxes into the stroma upon darkening deactivate carbon assimilation (*Hochmal et al., 2015*). Stromal pH and Mg^2+^ concentrations decrease upon darkening (*Ishijima et al., 2003*) and assist in enzyme regulation (*Minot et al., 1982*). These additional regulating parameters likely must be considered to fully simulate the CBC enzyme deactivation *in vivo*.

The results from this study strongly support the conclusion that the 2-CysPrx functions as Trx-oxidase *in vivo* which mediates the inactivation of Trx-dependent target proteins. This mechanism is needed to effectively inactivate or downregulate the assimilation pathways linked to the photosynthetic electron transport chain in darkness and upon a drop in photosynthetic active radiation. Beyond controlling the CBC and the malate valve, other metabolic processes such as diurnal starch accumulation and non-photochemical quenching are also affected by this pathway, and thus 2-CysPrx appears to be a global player in tuning the redox regulatory network of the chloroplast.

## Methods

### Growth of Arabidopsis

*A. thaliana* Col-0 wildtype and 2cysprxAB double insertion line (*Awad et al., 2015*) were grown on soil in 10 h light at 21°C and 14 h darkness at 18°C with 50% relative humidity. After stratification for 2 d in the dark at 4°C plants were grown at 80 μmol quanta m^-2^ s^-1^ (normal light: N) for isolating chloroplasts after 56 d and also for enzyme assays and NIR KLAS100 analysis after 28 d. Growth studies utilized a fluctuating light regime to mimic sun and shade in a natural environment and to explore growth advantages of plants with missing 2-CysPrx. To this end both lines were grown in N for 14 d or as indicated and then transferred to a home-built programmable LED-based light chamber with fluctuating light consisting of low light (L)-phases with 22 μmol quanta m^-2^s^-1^ and high light (H) phases with 800 μmol quanta m^-2^s^-1^. The fluctuating light consisted of 80 s L/10 s H or 40 s L/5 s H for 28d or as indicated. Control plants were kept in N. Diurnal light-dark cycles were as described above.

### Fluorescence of PSII, NIR-analysis of P700, PC and Fd absorption changes

Photosynthetic parameters were analyzed with a NIR-KLAS-100 spectrophotometer (Waltz, Germany, Effeltrich) using near infrared absorption spectroscopy (NIR) to visualize redox states of ferredoxin (Fd), plastocyanin (PC), photosystem I (PSI), and fluorescence from photosystem II (PSII) in leaves. Dark adapted plants were taken at the end of their normal dark phase and maintained in darkness. Two leaves were excised and sandwiched abaxially to fit into the 1×1 cm detection window for double-sided exposure. Fast kinetics was recorded in a 6 s time window with 1.5 s actinic light pulse of 162 μmol quanta m^-2^ s^-1^ to activate photosynthesis. Redox kinetics of the parameters was recorded for 4.5 s. Settings and device output were adjusted according to *Klughammer and Schreiber (2016)* and *Schreiber and Klughammer (2016)*.

### Gene cloning

The sequences encoding the mature proteins without targeting sequence of the Trx-f1, -m1, -m4, -x, CDSP32 and 2-CysPrxA from *A. thaliana* were amplified from leaf cDNA. Trx-f1, -m1, -m4, -x and CDSP32 were cloned into the *NdeI* and *BamHI* restriction sites of pET15b (Novagen). 2-CysPrxA was cloned into the *NdeI* and *EcoRI* restriction sites (underlined in the primers) of the pET28a (Novagen) using the primers described in Supplementary Table 9. Sequences were verified by DNA sequencing.

### Production and purification of recombinant proteins

The recombinant plasmids were introduced into the *E. coli* NiCo21 (DE3) strain (NEB). The bacteria were grown at 37°C, and protein production was induced by adding 100 μM isopropyl-β-D-thiogalactoside in the exponential phase. The bacteria were harvested by centrifugation at 5,000 rpm for 20 min and then resuspended in buffer A (50 mM phosphate buffer, pH 8.0, 10 mM imidazole, 250 mM NaCl) for Trx-f1, -m1, -m4, -x and CDSP32 and buffer B (50 mM phosphate buffer, pH 8.0, 10 mM imidazole, 250 mM NaCl, 40 μM of β-mercaptoethanol) for 2-CysPrxA. Cells were disrupted by sonication, and debris was sedimented at 13000 rpm for 30 min. The soluble fraction was loaded onto His-Select nickel affinity column (His-Select HF Nickel Affinity, ROTH) equilibrated with buffer A. Proteins were eluted with 50 mM phosphate buffer, pH 8.0, 250 mM imidazole, and 250 mM NaCl. Purified proteins were concentrated and dialyzed against 50 mM phosphate buffer, pH 8.0. Final purity was checked by 15% SDS-PAGE. Protein concentrations were determined spectrophotometrically using molar extinction coefficient at 280 nm of 17085 M^-1^ cm^-1^ for Trx-f1, 21095 M^-1^ cm^-1^ for Trx-m1, 19650 M^-1^ cm^-1^ for Trx-m4, 11585 M^-1^ cm^-1^ for Trx-x, 12170 M^-1^ cm^-1^ for CDSP32 and 20065 M^-1^ cm^-1^ for 2-CysPrxA. For oxidation, the recombinant 2-CysPrx was incubated in the presence of 1 mM H_2_O_2_ for 45 min and dialysed over night at 4°C with several changes of dialysis buffer (30 mM Tris-Cl, pH 8).

### Stroma isolation

Leaves were harvested from plants previously darkened for starch degradation and blended in homogenization buffer (300 mM sorbitol, 20 mM Tricine, 5 mM EGTA, 5 mM EDTA, 10 mM NaHCO_3_, 2 mM ascorbate and 1% bovine serum albumin; pH 8.4 with NaOH)(*Ströher et al.; 2008*). After filtering through 4 layers of gauze and nylon mesh the suspension was centrifuged, and the pellet re-suspended and spun again. Finally, chloroplasts were re-suspended in buffer containing 300 mM sorbitol, 30 mM KCl, 1 mM MgCl_2_, 0.2 mM KH_2_PO_4_, 2 mM ascorbate and 50 mM Hepes-NaOH pH 7.6. The suspension was stored at -80°C, thawed and centrifuged for 2 h to get stromal protein as supernatant.

### Enzyme activities in stromal extracts

The fructose 1,6-bisphosphatase (FBPase) activity was measured spectrophotometrically with the Shimadzu (UV-2401PC) at 340 nm (25°C). Stromal protein (100 μg) was preincubated in 30 mM Tris-HCl, pH 8.0, and 5 mM MgSO_4_ with DTT, with or without thioredoxin (Trx, 5 μM) as indicated. The preincubation solution (500 μl) was added to 500 μl reaction mix (0.1 mM NADP^+^, 30 mM Tris-HCl pH 8.0, 5 mM MgSO_4_, glucose-6-phosphate dehydrogenase and phosphoglucoisomerase (0.5 U each)) and oxidized 2-CysPrxA (final concentration: 5 μM 2-CysPrx_ox_). After recording the baseline for 3 min the assay was initiated by adding fructose 1,6-bisphosphate (0.6 mM FBP).

The malate dehydrogenase (MDH) test was conducted spectrophotometrically using the Cary 300 Bio UV-Visible (Varian, Middelburg, The Netherlands). The stromal protein (150 μg) was preincubated in buffer A (30 mM Tris-HCl, pH 8.0, 0.5 mM DTT, 0 or 10 μM Trx). Preincubation solution (500 μl) was mixed with 500 μl buffer B (30 mM Tris-HCl, pH 8.0, 0.4 mM NADPH and with or without 5 μM 2CysPrxAox). After recording the baseline at 340 nm, the assay was started by adding 2 mM oxaloacetic acid (OAA) as substrate.

### Enzyme activities in leaf extracts

For the MDH activity assay intact plants were exposed to high light of 650 μmol quanta m^-2^ s^-1^ for 30 min. After 0, 10, 30, 60 and 300 s in darkness, leaves were immediately frozen in liquid nitrogen. After addition of 450 μl 30 mM Tris-HCl, pH 8.0, to 100 mg pulverized leaf material, samples were vortexed for 20 s and centrifuged at 16,000 × g for 90 s. In order to measure the initial MDH-activity 200 μl of supernatant was added to a quartz cuvette containing 30 mM Tris-HCl, pH 8.0, 0.1 mM NADPH and 2 mM OAA. NADPH oxidation was measured at 340 nm (Cary 300) at 25 °C for 4 min. Total MDH-activity was determined after preincubation with 20 mM DTT for 30 min at RT. Background reactions were subtracted using a reference cuvette without OAA.

For PRK activity, pulverized leaf material (25 mg) from light-dark transitions was added to 450 μl PRK assay buffer (30 mM Tris-HCl pH 8.0, 1 mM EDTA, 40 mM KCl, 10 mM MgCl_2_) and processed as above. To measure the initial PRK activity 50 μl of supernatant was added to a quartz cuvette containing 2 mM ATP, 2.5 mM phospho*enol*pyruvate, 5 U/ml pyruvate kinase, 6 U/ml lactate dehydrogenase and 0.2 mM NADH. The reaction was started with 0.5 mM ribulose-5-phosphate and the activity was measured as a decrease in absorbance at 340 nm. The total PRK activity was quantified after preincubation with 20 mM DTT as above.

### FBPase redox state determination

100 μg of stroma was incubated under the conditions used for the FBPase activity assay. 500 μl of the reaction mixtures were stopped with one volume of 25% (w/v) trichloroacetic acid (TCA), and stored on ice for 1 hour. The mixtures were centrifuged at 13,000 rpm for 10 min and washed once with cold acetone. The pellet was resuspended in alkylation buffer 30 mM Tris-HCl pH 8.0 with 15 mM of mPEG_24_-mal (Methoxyl-PEG Maleimide) (Thermo Fischer Scientific) and incubated in the dark at room temperature for 1h. The alkylated mixtures were precipitated with 25% TCA, washed with cold acetone and resuspended in a loading buffer containing 30mM Tris-HCl pH8.0, 2.3 % SDS. The samples were separated in non-reducing SDS-PAGE (10 %), transferred onto nitrocellulose membrane and probed with an anti-FBPase antibody which was kindly provided by Pr. Jean-Pierre Jacquot (University of Lorraine, Nancy, France).

### Analysis of soluble and insoluble carbohydrates

Plants for carbohydrate analysis were harvested at day times as indicated (one hour before and after the light phase and in the middle of the day or night). Plants grew in short day with 10 h light with 80 μmol quanta m^-2^ s^-1^ at 21°C and 14 h darkness at 18 °C with 50% relative humidity. Carbohydrates were analyzed in 96-well plates according to *Leyva et al.(2008)* with minor modifications. Harvested WT and *2cysprxAB* leaves were grinded in liquid nitrogen and adjusted to 20-30 mg for optimal readings according to the linear calibration curves. After sedimentation at 13000 rpm for 10 min the sediment was washed with phosphate buffer and boiled at 100°C for 1 h in 3% hydrochloric acid to hydrolyze insoluble carbohydrates. After sedimentation at 13000 rpm for 10 min the supernatants were used as samples for insoluble carbohydrate determination. To 150 μl Anthron reagent (2 g l^-1^ in 75 % (v/v) sulfuric acid) 50 μl of sample or standard was added with gentle mixing prior to heating for 20 min at 100 °C. Insoluble acid-hydrolysable sugar was determined at 630 nm in three independent replicates using a glucose standard curve.

### Redox state of 2-CysPrx in high light to dark shifts

Plants were grown in growth light for 28 d as described before. WT was acclimated to 650 μmol quanta m^-2^ s^-1^ for 30 min (t = 0 s) and transferred to darkness (t = 5 to 120 s). Plant material was grinded and extracted with 50 mM Tris-HCl, pH 8.0 containing 100 mM N-ethylmaleimide (NEM) to alkylate free thiols while existing disulfides are kept oxidized. Separation on non-reducing 1D-SDS-PAGE allowed discrimination of labeled monomers (22 kDa) and oxidized intermolecular homodimers (44 kDa). The following Western Blot with luminescence output was used in combination with ImageJ software to quantify protein band intensities using 1 or 2 μg of protein extract for optimal exposure. Values given are percentage shares of reduced (NEM-labeled) and oxidized (oxidized SDS-PAGE) 2-CysPrx at each time point with n=4 replicates and ±SD.

### Kinetic modeling of the redox networks

Two models were implemented in Matlab. The first model was built to simulate the *in vitro* assay with pre-activation, Trx and FBPase and addition of oxidized 2-CysPrx at different concentrations. The parameters (concentrations, reaction constants) underlying the equations, the implementation of the redox potentials and the equations are given in *Supplementary Tables 1-8*. The second model was constructed to simulate the light-dark transition. In the presumed light phase, the fraction of reduced Fd was set to 50 %. To mimic the light-to-dark transition after 200 s, the Fd and FTR pool were turned fully oxidized, assuming the efficient drainage of electrons into the various Fd-dependent patways. The redox state of 2-CysPrx was clamped to 66% oxidized form and 34% reduced form according to the own date (*Figure 6B*).

### Statistical analysis

Statistics was assessed using one way analysis of variance (ANOVA) followed by post hoc analysis Tukey’s HSD (Honest significance difference) test (P≤0.05). Statistical results (significant difference) are indicated in the figures and represented by different letters. Statistical analyses were performed using IBM SPSS 20.0 software (IBM Corporation, Armonk, NY).

### Data availability

The authors declare that the data supporting the findings of this study are available within the article and the Supplementary Information files or are available upon reasonable request to the authors.

## Acknowledgements

The authors acknowledge support by the DFG (DI 346/14 & 17, SPP1710). We are grateful to Sergej Kakorin (Physical und Biophysical Chemistry, Bielefeld University) and Petra Lutter (Faculty of Biology, Bielefeld University) for help in kinetic modelling.

## Authors Contributions

MJV performed the in vitro analyses of FBPase and MDH, the Trx dependencies and also determined the FBPase redox state; KC prepared recombinant proteins, assisted in performing the *in vitro* experiments and their interpretation; WT measured MDH and PRK activity ex vivo, ML determined the 2-CysPrx redox state *ex vivo*, the carbohydrate levels, prepared recombinant proteins, and performed the NIR KLAS100 and the fluctuating light experiments; SMM performed the chlorophyll a fluorescence analyses of plate grown plants; MG constructed the kinetic model for simulating the regulation kinetics; KJD conceived the study, discussed the results and drafted the manuscripts. All authors contributed to data interpretation and writing.

## Conflict of Interest

The authors declare that there is no conflict of interest.

**Figure supplement 1. Thioredoxin-specificity of inactivation of FBPase by oxidized 2-CysPrx.**

**Figure supplement 2. Mathematical modeling and simulation of FBPase inactivation by 2-CysPrx_ox_ in the enzyme assay.**

**Figure supplement 3. Modeling and simulation of redox changes of FBPase and Trx-f1 upon darkening.**

**Table supplement 1. Values of variables used for modeling the enzyme assay.**

**Table supplement 2. Parameters used for modeling the enzyme assay as a reference.**

**Table supplement 3. Reaction equations describing the model of the enzyme test.**

**Table supplement 4. Rate expression for the three reactions of the enzyme test model.**

**Table supplement 5. Values of variables used for modeling the light-dark-transitions.**

**Table supplement 6. Parameters used for modeling the light-dark-transitions.**

**Table supplement 7. Reaction equations describing the model of the light-dark-transitions.**

**Table supplement 8. Rate expressions for the four reactions considered for the model of light-dark-transition**

